# Intergenerational effects of early adversity on survival in wild baboons

**DOI:** 10.1101/591248

**Authors:** Matthew N. Zipple, Elizabeth A. Archie, Jenny Tung, Jeanne Altmann, Susan C. Alberts

**Author notes:** Corresponding Author: Susan Alberts.

## Abstract

In humans and nonhuman animals, early life adversity can affect an individual’s health, survival, and fertility for many years after the adverse experience. However, whether early life adversity also imposes intergenerational effects on the exposed individual’s offspring is not well understood. Here, we fill this gap by leveraging prospective, longitudinal data on a wild, long-lived primate. We find that juveniles whose mothers experienced early life adversity exhibit high mortality before age 4, and this effect is independent of the juvenile’s own experience of early adversity. Furthermore, our results point towards a strong role for classic parental effects in driving these effects: mothers that experienced early life adversity displayed reduced viability in adulthood, which in turn led to reductions in offspring survival. Importantly, these mothers’ juvenile offspring often preceded them in death by 1 to 2 years, indicating that, for high adversity mothers, the quality of maternal care declines near the end of life. While we cannot exclude direct effects of a parent’s environment on offspring quality (e.g., transgenerational epigenetic changes), our results are most consistent with a classic parental effect, in which the environment experienced by a parent affects its future phenotype and therefore its offspring’s phenotype. Together, our findings demonstrate that adversity experienced by individuals in one generation can have strong effects on the survival of offspring in the next generation, even if those offspring did not themselves experience early adversity.

---

An individual’s health, survival, and fertility can be profoundly shaped by its early life environment (1). For example, in humans, low early life socioeconomic status predicts increased risk of coronary heart disease (2–4), stroke (2, 5, 6), type II diabetes (7), poor perceived health (8), and all-cause mortality (9, 10) in adulthood. Similarly, numerous studies of wild mammals (11–14) and birds (15–17) find that adult fecundity is reduced in animals that experienced adverse early life environments, and a few have also found an effect of early life adversity on adult survival (13–15, 18).

If the effects of early adversity extend to the descendants of exposed individuals, the epidemiological and evolutionary impact of these effects would be further amplified. However, evidence from humans for intergenerational effects that result directly from the early life experience of the parent is mixed, as studies have produced somewhat contradictory results (19–22). For example, a study of the Överkalix population in Sweden identified strong, contrasting effects of grandparents’ exposure to early-life food scarcity on grand-offspring survival, depending on small differences in the age at which the grandparent was exposed to scarcity (22). Similarly, two studies of the same population exposed *in utero* to the Dutch hunger winter (a well-studied famine that resulted from a German blockade of the Netherlands during the winter of 1944-1945) found contradictory, sex-specific intergenerational effects, in one case suggesting an intergenerational effect that depended only upon the mother’s early experience (20), and in the other case an effect that depended only upon the father’s early experience (19). Furthermore, any possible effects of parental or grandparental adversity on future generations are assumed to be transgenerational, operating as a result of inherited epigenetic changes (19–22). Yet no genetic validation of this assumption has been carried out, and it remains possible that a simpler intergenerational pathway explains such results. Specifically, early adversity experienced by a parent may act as a classic parental effect by changing the parent’s phenotype, which in turn influences the offspring’s phenotype (23–26).

The best evidence for intergenerational effects of early adversity comes from several laboratory studies of short-lived animals, which find strong relationships between a female’s early life environment and the body size of her offspring [(27–36), reviewed in (37), but see (38) for a rare example in wild house wrens]. These findings provide important proof-of-principle that intergenerational effects of early adversity can occur. However, these studies do not address whether intergenerational effects of early adversity occur in natural populations of long-lived animals. And while a few studies of captive animals have demonstrated a relationship between a female’s early environment and her offspring’s survival or reproduction (39–41), the ecological validity of these findings has yet to be verified by studying intergenerational fitness effects in a population of wild and/or long-lived animals. By working in a natural population, we are able to guarantee that animals are exposed only to natural levels of early adversity, and are also subject to any social factors which might mitigate or aggravate the influence of those early adverse events.

Addressing whether the effects of early adversity in one generation affect reproduction or survival in the next is challenging because of the difficulties of linking high-quality data on early adversity in one generation to health and survival outcomes in the next. Here, we overcome these challenges by taking advantage of a prospective longitudinal dataset from a natural primate population: the baboons of the Amboseli ecosystem in southern Kenya (42). This dataset includes 45 years of individual-based data on early adversity, and real-time observations of later-life survival outcomes for hundreds of subjects with known maternities and grand maternities. Moreover, unlike many human populations, we do not observe inter-generational transmission of adverse conditions; that is, offspring of females who experienced early life adversity are not more likely to experience early life adversity themselves—allowing us to avoid this common confound in human societies.

To test for intergenerational effects of early adversity, we focused on early adversity experienced by female baboons who later became mothers, and whose offspring were also in our data set. We asked whether the early adversity experienced by these females (“maternal early adversity”) predicted the survival of their juvenile offspring in the next generation, after controlling for the early adversity directly experienced by the offspring themselves.

We considered five types of early adverse conditions (Table 1), based on previous work in our study population that demonstrated effects of these conditions on a female baboon’s own adult survival (18). These included: (i) maternal death during development (0-4 years of age), which indicates the loss of an important source of social support, physical protection, and nutrition (43, 44), (ii) being born to a low-ranking mother, which influences growth rates and age at maturation (45–47) (iii) being born into a large social group (and thus experiencing high density conditions and high levels of within-group competition) (11, 45, 48) (iv) being born during a drought, which reduces fertility in adulthood (11, 49), and (v) experiencing the birth of a close-in-age younger sibling, which may reduce maternal investment received during development (50). Importantly—and in contrast to human studies (51)—sources of early adversity are not strongly correlated in our population (Table S1).

**Table 1.**
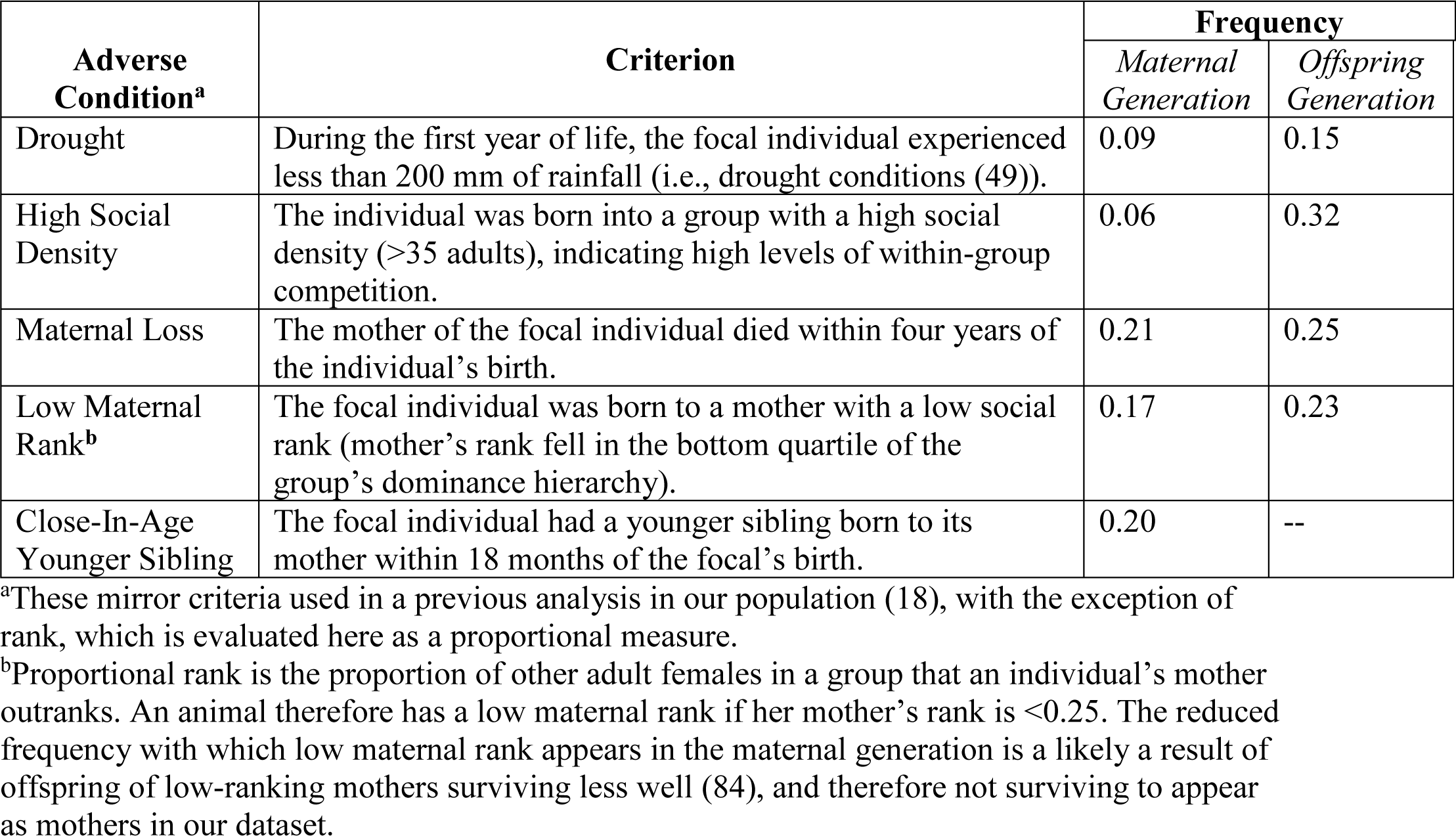
Early adverse conditions and the frequencies with which they occur in maternal and offspring generations of our dataset.

| Adverse Condition<sup>a</sup> | Criterion | Frequency |  |
| --- | --- | --- | --- |
|  |  | Maternal Generation | Offspring Generation |
| Drought | During the first year of life, the focal individual experienced less than 200 mm of rainfall (i.e., drought conditions (49)). | 0.09 | 0.15 |
| High Social Density | The individual was born into a group with a high social density (>35 adults), indicating high levels of within-group competition. | 0.06 | 0.32 |
| Maternal Loss | The mother of the focal individual died within four years of the individual's birth. | 0.21 | 0.25 |
| Low Maternal Rank<sup>b</sup> | The focal individual was born to a mother with a low social rank (mother's rank fell in the bottom quartile of the group's dominance hierarchy). | 0.17 | 0.23 |
| Close-In-Age Younger Sibling | The focal individual had a younger sibling born to its mother within 18 months of the focal's birth. | 0.20 | -- |

<sup>a</sup>These mirror criteria used in a previous analysis in our population (18), with the exception of rank, which is evaluated here as a proportional measure.
<sup>b</sup>Proportional rank is the proportion of other adult females in a group that an individual's mother outranks. An animal therefore has a low maternal rank if her mother's rank is <0.25. The reduced frequency with which low maternal rank appears in the maternal generation is a likely a result of offspring of low-ranking mothers surviving less well (84), and therefore not surviving to appear as mothers in our dataset.

## Results

We built a mixed effects Cox proportional hazards model of offspring survival during the juvenile period that included early adversity measures present in the mother’s and the offspring’s early life as binary fixed effects. We defined the juvenile period based on survival until age 4, near the age of menarche for females and earliest dispersal for males in this population (52). We included data on maternal early adversity for all five adverse early life conditions, and we included data on offspring early adversity for four of the five conditions. We excluded the birth of a close-in-age younger sibling for the offspring generation because the survival of the focal offspring strongly affects the length of the subsequent birth interval (i.e., offspring that die shortly after birth also have the closest-in-age younger siblings). We included maternal and grandmaternal ID as random effects. In total, we used data collected from 1973-2017 to analyze the survival of 687 offspring (46.5% males) born to 169 females (mean 4.1 offspring per female, range 1-12) for whom we had data on all five adverse conditions in the mother’s early life, and all four adverse conditions in the offspring’s early life.

Each adverse condition was scored as present or absent for each subject, and each one affected a minority of our study subjects (range 6%-34%). Mothers and offspring had similar chances of experiencing adverse conditions, except for social density: offspring were more likely than mothers to be born into large social groups because of population growth over the 5-decade study period (Table 1). Unlike typical patterns of early adversity in human populations (51), sources of early life adversity in our population were not strongly correlated: with the exception of maternal rank in the mother’s and offspring’s generations (p<0.0001, r=0.40), no adverse condition explained more than 4% of the variance in any other condition, either within or between generations (Table S1).

### Maternal Early Life Adversity and Offspring Survival

Our full multivariate Cox proportional hazards model for offspring survival (Table S2) included all 9 early adverse conditions (five for mothers and four for offspring). We found strong negative effects of two characteristics of the *mother’s* early life environment on their offspring’s survival during the first 4 years of life: maternal loss (hazard ratio = 1.48, p=0.006) and presence of a close-in-age younger sibling (HR = 1.39, p=0.03). Following backwards model selection, these two characteristics remained the only significant maternal early life predictors of offspring survival (Table 2, Figure 1, along with two conditions in the offspring’s early life environment: see below). Specifically, offspring whose mothers experienced early maternal loss experienced a 48% higher probability of dying throughout the first four years of life than unaffected offspring, and offspring whose mothers had a close-in-age sibling experienced a 39% higher probability of dying than unaffected offspring. This effect is striking especially considering that a median of 7.0 and 8.0 years separated the offspring’s own birth from the mother’s experience of maternal loss or birth of a close-in-age sibling, respectively. Notably, previous work in our population found that these two sources of adversity—maternal loss and the presence of a close-in-age younger sibling during early life—are also sources of mortality risk once females reach adulthood, and in fact are the two strongest predictors of adult survival among six different early-life conditions considered (18). Hence, early-life conditions that are especially adverse for females when they reach adulthood also negatively affect the survival of their offspring.

**Table 2.** Reduced model of the effects of maternal and offspring early adversity on offspring survival during early life (R^2^ =0.07).

| Generation | Parameter<sup>a</sup> | Coefficient | Hazard Ratio<br>(95% CI) | p value | Interpretation |
| --- | --- | --- | --- | --- | --- |
| *Maternal* | Maternal Loss | 0.37 | 1.44<br>(1.10-1.90) | **0.009** | Offspring survive less well if their mother experienced maternal loss during her early life. |
|  | Close-in-age Younger Sibling | 0.35 | 1.42<br>(1.06-1.90) | **0.018** | Offspring survive less well if their mother had a close in age younger sibling during her early life. |
| *Offspring* | Maternal Loss | 0.68 | 1.98<br>(1.53-2.56) | **$3 \times 10^{-7}$** | Offspring survive less well if they experienced maternal loss within four years of their birth. |
|  | Low Maternal Rank | 0.42 | 1.54<br>(1.17-2.01) | **0.002** | Offspring survive less well if they were born to a low-ranking mother. |

<sup>a</sup>An alternative model that considered cumulative maternal adversity was not a better or worse fit than the reduced multivariate maternal adversity model (see Table S7. For both the model presented here and that in Table S7, $R^2=0.07$ , log likelihood = -1598).

**Figure 1.**
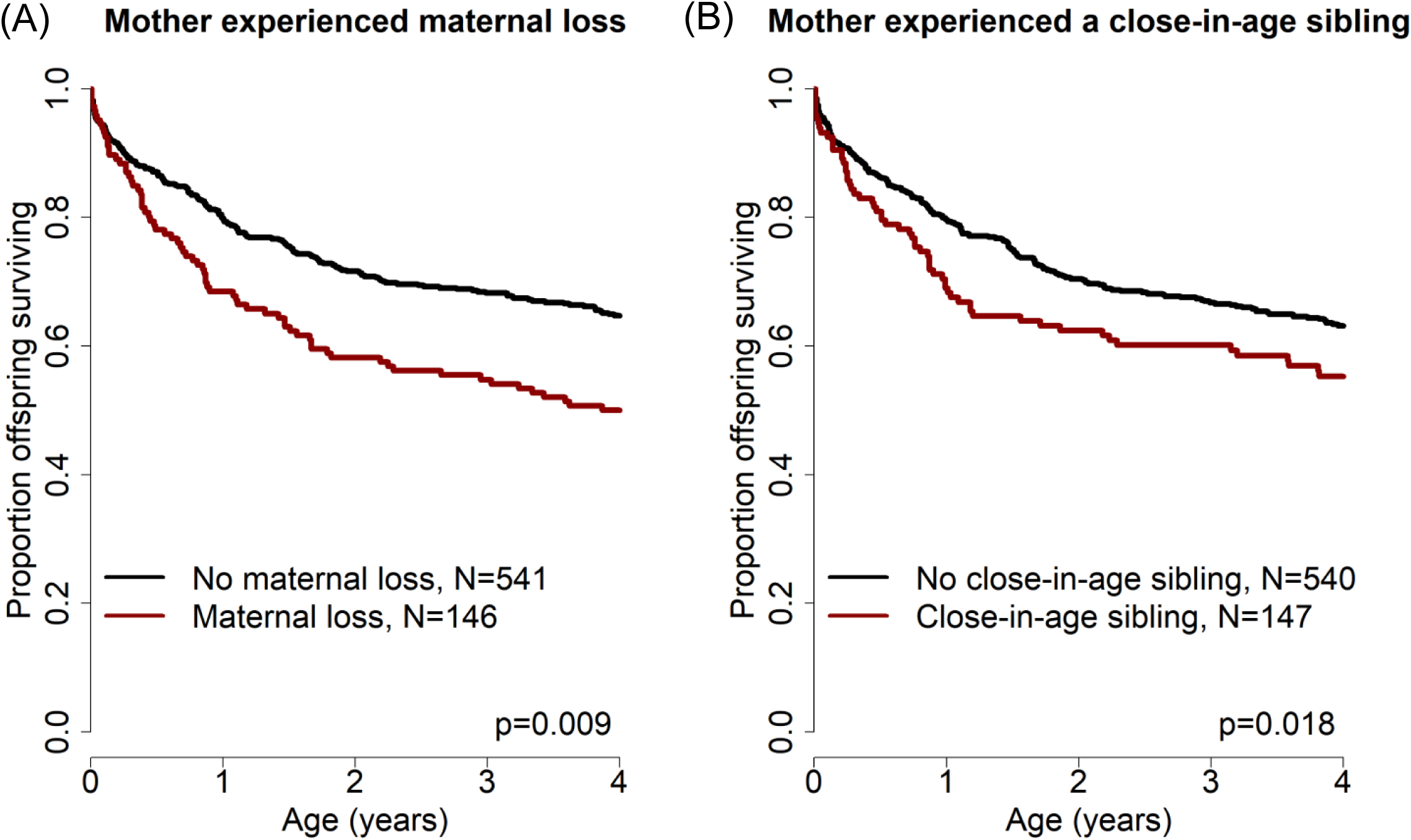
Offspring survival was influenced by characteristics of their mothers’ early-life environments. Offspring survived relatively less well during the juvenile period if (A) their mother lost her own mother during her early life and/or (B) their mother experienced a close-in-age younger sibling.

Both the full and reduced models of offspring survival also included two conditions in the *offspring’s* early life environment as significant predictors of juvenile survival. Specifically, maternal loss experienced by the offspring and low maternal rank during the offspring’s juvenile period had strong negative effects on offspring survival (Table 2, maternal death: HR = 1.95, p=5×10^−7^, low maternal rank: HR=1.43, p=0.025). Thus, maternal loss in the offspring’s generation had a stronger effect on offspring survival (nearly doubling offspring mortality risk) than maternal loss in the mother’s generation. In contrast, the effect of having a low-ranking mother, which was associated with a 43% increase in offspring mortality risk, was comparable in its effect size to the two significant predictors from the maternal generation (maternal loss and close-in-age sibling for the mother, 48% and 39% increase in offspring mortality, respectively). Thus, two adverse conditions in a mother’s early life had as large or larger of an impact on her offspring’s survival than all but one adverse condition experienced by the offspring directly.

### Maternal Viability and Offspring Survival

The strong effect of the mother’s death on the survival of offspring to age four years (Table 2) could arise if offspring die *after* their mothers die: even after weaning (approximately 1.5 years of age), juvenile baboons rely on their mothers for social support and social learning (53). Alternatively, these offspring may die *before* their mothers die if those mothers are themselves in poor condition. To distinguish these alternatives, we modeled offspring survival to age 2 years (halfway through the juvenile period) as a function of maternal death during years 2-4 after an offspring’s birth (i.e., the two years that *followed* the offspring survival period modeled in the response variable). In this analysis we considered only the subset of offspring in our dataset whose mothers survived the entirety of the first two years of the offspring’s life, and for whom we were able to evaluate the four significant predictors of offspring survival identified above and in Table 2 (N=671). Our results showed that offspring were less likely to survive during the first two years of life if they were born to mothers who died 2-4 years after their birth. In other words, these offspring were more likely to die even when their mother was still alive (hazard ratio=1.50 [1.01-2.23], p=0.045).

To test whether this link between offspring survival and maternal viability was driven by maternal early adversity, we next partitioned our analysis of offspring survival to age 2 based on whether the mother experienced either maternal loss or a close-in-age younger sibling (i.e., either or both of the two maternal early life conditions that significantly predicted their offspring’s survival; Table 2). We found that, among offspring whose mothers experienced either or both of these two conditions (N=247), maternal death in years 2-4 after the offspring’s birth significantly predicted reduced offspring survival to age 2 years (Figure 2a, hazards ratio=1.78, 95% CI = [1.05-3.01], p=0.03). Maternal death in the same period did not, however, predict reduced offspring survival when mothers had not experienced maternal loss or a close-in-age younger sibling (N=424; Figure 2b, hazard ratio=1.21, 95% CI = [0.7-2.2], p=0.53). This finding is consistent with the hypothesis that maternal early life adversity results in low maternal viability in adulthood, which in turn results in both earlier death for adult females and a reduction in their ability to successfully raise offspring towards the end of their lives.

**Figure 2.**
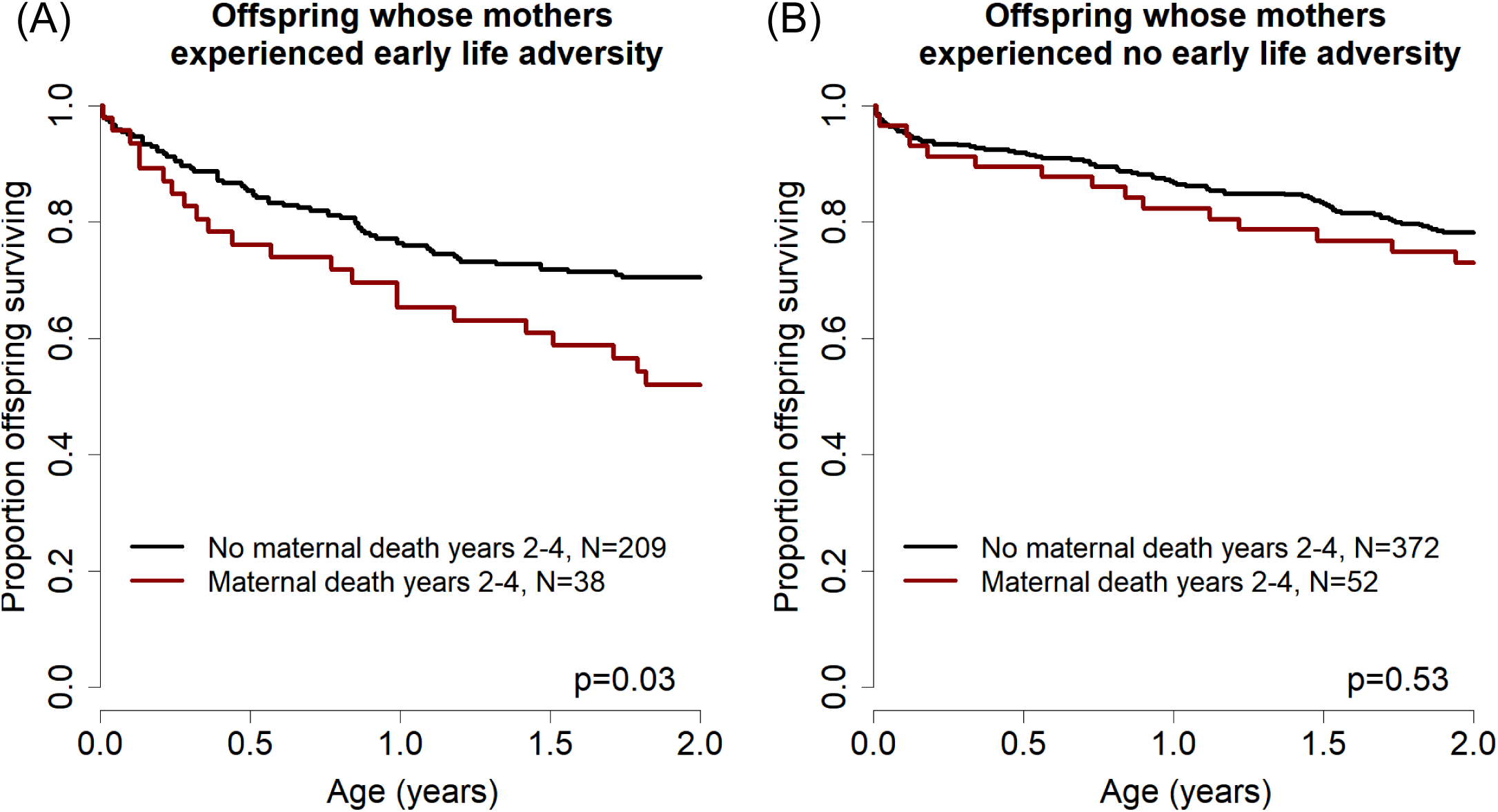
Effects of maternal adversity on offspring survival are explained by reduced maternal viability. (A) Among those offspring whose mothers experienced significant early life adversity (maternal loss and/or a competing younger sibling), maternal death in years 2-4 was associated with poor offspring survival in years 0-2 after birth, while the mother was still alive. (B) In contrast, among those offspring whose mothers did not experience early life adversity, maternal death in years 2-4 was not associated with reduced offspring survival during the first 2 years of life.

### Maternal Early Life Adversity and Quantity of Maternal Care

We hypothesized that developmental constraints imposed on females by early life adversity could lead to reduced survival in their offspring as a result of two non-mutually exclusive mechanisms. First, mothers that experienced early life adversity might provide lower levels of maternal care to their offspring than other mothers, through differences in either maternal behavior or physiology (e.g. reduced nutrient content in milk). Second, females who experienced early adversity might also exhibit reduced egg and/or amniotic environmental quality (an established mechanism for transmission of maternal effects (37, 54)). We were able to partially test the first hypothesis by drawing on longitudinal behavioral data for this population. Specifically, we built linear mixed effects models to test whether maternal early adversity affected the proportion of time during 10-minute focal follows that a mother spent either carrying or suckling her dependent infants. Fixed effects in the model included maternal viability (a binary variable indicating whether the mother survived for four years after offspring birth), the two early adverse circumstances experienced by the mother that affected offspring survival (the mother’s maternal loss and close-in-age sibling), offspring age (both as a linear and as a quadratic term), maternal rank, maternal age, the number of adult females in the social group, and season. We also included maternal ID, offspring ID, group ID, and observer ID as random effects.

Neither maternal viability nor either of the two maternal early adversity effects was a significant predictor of the proportion of time that mothers spent carrying or suckling their infants in either full multivariate models or models resulting from backwards selection (p>0.1 in all cases, Tables S3-S6). However, the proportion of time that mothers spend carrying and suckling offspring are relatively coarse metrics of maternal care, and it could be that more fine-grained measures of maternal care (which we do not regularly collect) could reveal a relationship between maternal early adversity and future maternal care.

## Discussion

We have demonstrated that adverse environmental conditions during the early life of a female baboon, which are already known to negatively affect both her survival (18) and her reproduction (11) in adulthood, also reduce the survival of her offspring. Importantly, this effect is independent of the environment experienced by those offspring themselves (Figure 1). The reduction in offspring survival is likely linked to reductions in maternal viability: mothers that experienced early life adversity are significantly less able to successfully raise offspring born near the ends of their lives, while the same is not true for mothers that did not experience early life adversity (Figure 2). Together, these findings support the hypothesis that early life adversity produces constraints during development that lead not only to reduced adult survival and lifetime reproductive success (18) but also to a reduced ability to successfully raise those offspring that are produced.

The results reported here help to fill a key gap in the literature concerning the intergenerational effects of early life adversity on survival. Results from human studies have yielded inconsistent results on this topic thus far: different studies on the same populations have reported contradictory sex-specific effects on health (19, 20) or have found that small differences in the age at which probands’ parents or grandparents were exposed to adversity can lead to a reversal in the direction of these effects (21, 22). Among studies in non-human animals, several studies in fish (55, 56), reptiles (57), birds (58, 59), and ungulates (60–64) have found that parental body condition at the time of offspring birth influences offspring survival, but none have linked parents’ early adverse experiences to offspring survival. Additionally, while previous studies have identified effects of parental early adversity on offspring traits in a limited number of systems (37, 39, 40), ours is the first to link parental early adversity to offspring fitness outcomes in a wild, long-lived animal.

Our findings help to explain the persistence of health deficits across generations (65–67), by revealing that in long-lived primates, the early life experiences of mothers have important implications for offspring health and survival. Recent studies in humans have demonstrated that conditions experienced by mothers during pregnancy (e.g., low SES, psychosocial stress, mood dysregulation, prenatal smoking) can affect HPA axis regulation (68, 69) and birthweight (65, 66) in her offspring. These and other maternal characteristics present during pregnancy are influenced not only by mothers’ experiences in adulthood, but also by the long-term effects of environmental conditions experienced in mothers’ early lives (54, 66). Our findings therefore motivate future work to test for comparable intergenerational fitness effects of early adversity in humans and other non-human animals.

Our findings are consistent with the hypothesis that early adversity results in intergenerational effects of developmental constraints (11, 70–72) and are not consistent with an intergenerational predictive adaptive response hypothesis (71, 73, 74). Rather than being buffered against the effects of maternal loss, those offspring that experienced maternal loss and whose mothers had also experienced maternal loss were more likely, not less likely, to die, as compared to offspring that experienced maternal loss but whose mothers did not. Thus, offspring experience constraints not only as a result of their own early environment, but also as a result of their mother’s developmental history, including events that occurred years before the offspring’s own conception. Our results are consistent with the hypothesis that a female’s condition at the time of her offspring’s conception and/or birth reflects her previous experiences, and that her condition thereby influences the development and survival of her offspring (54, 75, 76).

Finally, our study provides insight into how the effects of early adversity may be transmitted from parent to offspring. Intergenerational transmission of adversity has often been viewed as a potential transgenerational effect – i.e., environmental exposures in one generation that directly alter the biology of animals born at least one generation later, perhaps via the inheritance of environmentally-induced epigenetic changes (77). While we cannot exclude this possibility, our results suggest that a simpler mechanism may be operating. The intergenerational effects of early adversity on offspring survival that we present here are consistent with a classic parental effect (19-22) in which early life adversity affects the phenotypic quality of the mother during adulthood, and her resulting deficits directly affect her offspring’s development. This mode of transmission – in which intergenerational transmission of parental early adversity operates via differences in parental phenotype, rather than via transgenerational effects that determine offspring phenotypes – may be more widespread than is typically thought (78). This insight may have important implications for understanding the persistence of human health deficits across generations and inform the approaches taken to intervene in this transmission (79).

## Methods

### Study system

The Amboseli Baboon Research Project is a long-term longitudinal study of wild baboons living in and around Amboseli National Park, Kenya. A detailed description of the study system can be found elsewhere (42). Researchers have continuously collected behavioral, environmental, and demographic data from the population since 1971. All subjects are visually recognized, and near-daily censuses allow us to precisely document the timing of demographic events, including the birth and death of study individuals. Critical to this study, we have continuously collected near-daily measures of group size, daily rainfall levels, and monthly calculations of social dominance rank (80), and we can accurately assign the dates of birth of all mothers and offspring born into study groups as well as the dates of all juvenile and adult female deaths.

### Study Subjects

In our analyses of offspring survival, we included individuals who met two criteria: (i) we were able to evaluate each of the five sources of maternal early life adversity and four sources of offspring early life adversity outlined below; and (ii) they lived in social groups that fed exclusively on wild foods rather than having their diet supplemented with human-sourced refuse. Although transmission of paternal early adversity may also occur in our population, we did not consider it here because we knew paternal identities for only a subset of our study subjects and had early life data on only a limited number of fathers. Our analysis ultimately relied on data spanning more than four decades, from 1973 to 2017.

### Measuring Early Life Adversity

Previous work in the Amboseli population defined six binary indicators of early life adversity and considered a single index of cumulative adversity based on the sum of these indicators (18). This cumulative adversity index is a strong predictor of adult lifespan: females that experienced high levels of early life adversity (i.e., a greater number of adverse early life conditions) but still survived to adulthood lived dramatically shorter lives compared to females that did not experience adversity (18). In addition to the five sources of early adversity discussed above, this previous analysis also considered early social connectedness (social integration versus social isolation) as a sixth source of adversity (18). Social connectedness data are missing for some mothers who were born relatively early in the long-term study. To maximize our sample size, we therefore did not include measures of social connectedness in this analysis. Our operational definitions for each source of adversity mirrored those used by Tung *et al* (18) for the remaining five conditions, except that here we employed measures of proportional rather than ordinal dominance rank (i.e., rank measured as a proportion of females that the focal individual dominates, rather than her ordinal rank number). We also built an index of cumulative maternal adversity, but because that model did not fit the data better than our reduced multivariate model (in contrast to the results for adult female survival (18)) we report the multivariate model in the main text. The alternative model based on cumulative maternal adversity is presented in Table S7.

### Statistical Analysis

We built a mixed effects Cox proportional hazards model of offspring survival during the first four years of life using the R package coxme (81, 82). The response variable in our model was the age at which offspring death occurred (if at all) during the first 4 years of life. We considered offspring survival to age 4 as the key survival period of interest because it roughly corresponds to the end of the juvenile period for baboons (52). Offspring that survived beyond age 4 were treated as censored individuals who survived until at least age 4. In our models of offspring survival as a function of maternal viability (Figure 2), we altered the first model to predict survival during the first two years of life as a function of maternal survival during years 2-4 after offspring birth. To test for effects of maternal adversity on the quantity of maternal care we built linear mixed effects models using the R package lme4 (83) (see Table S8 for model syntax).

