## Supplementary material for "Intergenerational effects of early adversity on survival in wild baboons": Tables S1-S8

Table S1. Pearson correlation coefficients between binarized sources of early life adversity[a]

| r (adj. p: Holm's method) | Maternal Loss (Mom) | Maternal Loss (Offsp) | Low Maternal Rank (Mom) | Low Maternal Rank (Offsp) | Drought (Mom) | Drought (Offsp) | Large Group (Mom) | Large Group (Offsp) | Close-in-age Sibling (Mom) |
|---|---|---|---|---|---|---|---|---|---|
| Maternal Loss (Mom) | --- | | | | | | | | |
| Maternal Loss (Offsp) | 0.12[b]<br>0.07[d] | --- | | | | | | | |
| Low Maternal Rank (Mom) | -0.07<br>1.00 | 0.07<br>1.00 | --- | | | | | | |
| Low Maternal Rank (Offsp) | 0.07<br>1.00 | 0.06<br>1.00 | **0.40**<br>**<0.0001** | --- | | | | | |
| Drought (Mom) | 0.06<br>1.00 | 0.06<br>1.00 | -0.02<br>1.00 | 0.04<br>1.00 | --- | | | | |
| Drought (Offsp) | 0.01<br>1.00 | 0.05<br>1.00 | -0.02<br>1.00 | 0.04<br>1.00 | -0.06<br>1.00 | --- | | | |
| Large Group (Mom) | 0.13<br>**0.01** | 0.09<br>0.58 | -0.07<br>1.00 | 0.06<br>1.00 | -0.02<br>1.00 | -0.06<br>1.00 | --- | | |
| Large Group (Offsp) | -0.06<br>1.00 | 0.01<br>1.00 | -0.12<br>0.041 | -0.02<br>1.00 | -0.10<br>0.27 | 0.17<br>**0.0001** | 0.03<br>1.00 | --- | |
| Close-in-Age Sibling (Mom) | -0.08<br>0.89 | 0.05<br>1.00 | 0.09<br>0.37 | -0.17<br>**0.0003** | -0.11<br>0.075 | 0.03<br>1.00 | -0.12<br>0.06 | 0.08<br>1.00 | --- |

[a]Correlations are calculated for binary measures of adversity within and between generations

[b]Pearson's correlation coefficient (r), reported for each pair of variables. Coefficients > 0.2 appear in bold

[c]p-value for r, reported for each pair of variables

[d]p-value following Holm's method for correcting for multiple testing

Table S2. Full mixed effects Cox proportional hazards model:

| Generation | Parameter | Coefficient[a] | Hazard Ratio (95% CI) | p value | Interpretation |
|---|---|---|---|---|---|
| *Maternal* | Maternal Loss | 0.39 | 1.47 (1.12-1.95) | **0.006** | Offspring survived less well if their mother experienced maternal loss during her early life. |
| | Close-In-Age Younger Sibling | 0.33 | 1.39 (1.03-1.89) | **0.03** | Offspring survived less well if their mother had a close-in-age younger sibling during her early life. |
| | Low Maternal Rank | 0.22 | 1.25 (0.89-1.76) | 0.19 | |
| | Drought | 0.14 | 1.15 (0.77-1.71) | 0.50 | |
| | High Social Density | -0.12 | 0.89 (0.52-1.51) | 0.66 | |
| *Offspring* | Maternal Loss | 0.67 | 1.95 (1.51-2.54) | **$5x10^{-7}$** | Offspring survived less well if their mother died within four years of their birth. |
| | Low Maternal Rank | 0.35 | 1.43 (1.05-1.94) | **0.03** | Offspring survived less if well if they were born to a low-ranking mother. |
| | Drought | -0.29 | 0.75 (0.52-1.08) | 0.12 | |
| | High Social Density | -0.07 | 0.93 (0.71-1.22) | 0.61 | |

[a]In all cases, positive coefficients indicate a higher hazard ratio in the presence of the adverse condition

Table S3. Full mixed effects model of proportion time spent carrying offspring

| Parameter | Coefficient | Standard Error | p value | Interpretation |
|---|---|---|---|---|
| Intercept | 0.955 | 0.028 | | |
| Offspring Age (Weeks) | -0.034 | 0.0007 | **<0.0001** | Proportion of time that a mother carries her infant declines in a quadratic fashion as infant age increases (concave up). |
| Offspring Age (Weeks) Squared | 0.0003 | 0.00001 | **<0.0001** | |
| Season (Wet) | 0.030 | 0.006 | **<0.0001** | Proportion of time that a mother carries her infant is higher in the wet season. |
| Number of Adult Females | 0.0008 | 0.0009 | 0.39 | |
| Maternal Viability | 0.018 | 0.013 | 0.16 | |
| Maternal Age (Years) | 0.0003 | 0.001 | 0.82 | |
| Mother is Low Ranking | 0.011 | 0.013 | 0.42 | |
| Mother Experienced Maternal Loss | 0.009 | 0.015 | 0.56 | |
| Mother Experienced Close-In-Age Younger Sibling | 0.012 | 0.016 | 0.43 | |

Table S4. Reduced mixed effects model of proportion time spent carrying offspring

| Parameter | Coefficient | Standard Error | p value | Interpretation |
|---|---|---|---|---|
| Intercept | 0.982 | 0.028 | | |
| Offspring Age (Weeks) | -0.034 | 0.0007 | **<0.0001** | Proportion of time that a mother carries her infant declines in a quadratic fashion as infant age increases (concave up). |
| Offspring Age (Weeks) Squared | 0.0003 | 0.00001 | **<0.0001** | |
| Season (Wet) | 0.031 | 0.006 | **<0.0001** | Proportion of time that a mother carries her infant is higher in the wet season. |

Table S5. Full mixed effects model of proportion time spent nursing offspring

| **Parameter** | **Coefficient** | **Standard Error** | **p value** | **Interpretation** |
|---|---|---|---|---|
| Intercept | 0.875 | 0.025 | | |
| Offspring Age (Weeks) | -0.039 | 0.0007 | <**0.0001** | Proportion of time that a mother nurses her infant declines in a quadratic fashion as infant age increases (concave up). |
| Offspring Age (Weeks) Squared | 0.0005 | 0.00001 | <**0.0001** | |
| Season (Wet) | 0.013 | 0.006 | **0.03** | Proportion of time that a mother nurses her infant is higher in the wet season. |
| Number of Adult Females | -0.001 | 0.0009 | 0.19 | |
| Maternal Viability | 0.018 | 0.011 | 0.12 | |
| Maternal Age (Years) | -0.0007 | 0.001 | 0.53 | |
| Mother is Low Ranking | 0.021 | 0.012 | 0.07 | |
| Mother Experienced Maternal Loss | 0.009 | 0.014 | 0.53 | |
| Mother Experienced Close-In-Age Younger Sibling | 0.006 | 0.014 | 0.66 | |

Table S6. Reduced mixed effects model of proportion time spent nursing offspring

| **Parameter** | **Coefficient** | **Standard Error** | **p value** | **Interpretation** |
|---|---|---|---|---|
| Intercept | 0.875 | 0.025 | | |
| Offspring Age (Weeks) | -0.039 | 0.0007 | <**0.0001** | Proportion of time that a mother nurses her infant declines in a quadratic fashion as infant age increases (concave up). |
| Offspring Age (Weeks) Squared | 0.0005 | 0.00001 | <**0.0001** | |
| Season (Wet) | 0.013 | 0.006 | **0.03** | Proportion of time that a mother nurses her infant is higher in the wet season. |

Table S7. Alternative mixed effects survival model that includes cumulative maternal adversity instead of multivariate adversity conditions ($R^2$=0.07, log likelihood = -1598)

| Generation | Parameter | Coefficient | Hazard Ratio (95% CI) | p value | Interpretation |
|---|---|---|---|---|---|
| Maternal | Cumulative Maternal Adversity | 0.27 | 1.31 (1.12-1.52) | **0.0007** | Offspring survived less well if their mother experienced more adversity during her early life. |
| Offspring | Maternal Loss | 0.66 | 1.94 (1.50-2.52) | **$6x10^{-7}$** | Offspring survived less well if their mother died within four years of their birth. |
| | Low Maternal Rank | 0.30 | 1.36 (1.03-1.78) | **0.03** | Offspring survived less if well if they were born to a low ranking mother. |

Table S8. Final reduced models written out in R syntax.

<table>
<tr><td>Adversity Model:</td></tr>
<tr><td>Surv(Age_4, Death_4)~ML+MS+OL+OR+(1|Maternal ID)+(1|Grandmaternal ID)</td></tr>
<tr><td></td></tr>
<tr><td>Maternal Loss Model:</td></tr>
<tr><td>Surv(Age_2, Death_2)~ML+(1|Maternal ID)</td></tr>
<tr><td></td></tr>
<tr><td>Carrying Model:</td></tr>
<tr><td>prop_carrying~Age+Age^2+Season+(1|Maternal ID)</td></tr>
<tr><td></td></tr>
<tr><td>Suckling Model:</td></tr>
<tr><td>prop_suckling~Age+Age^2+Season+(1|Maternal ID)</td></tr>
</table>

ML: Maternal loss in the mother's generation
MS: Maternal exposure to a close-in-age younger sibling
OL: Maternal loss in the offspring's generation
OR: Low maternal rank in the offspring's generation
Age_4/Age_2: Offspring age at death/censor. Maximum age = 4 or 2 depending on analysis
Death_4/Death_2: Binary indicator of whether death occurred at the offspring's listed age. (1= death, 0 = censored)
prop_carrying/prop_suckling: Proportion of time that a mother spent carrying/nursing her infant during the focal sample
Age/Age^2: Offspring age at the time of focal sample
Season: Binary; Wet (November-May) or dry (June-October)
